# Intestinal Enteroid Monolayers Model the Human Intestinal Environment for *Escherichia coli* Infection

**DOI:** 10.1101/2021.12.20.473596

**Authors:** Jason Small, Alison Weiss

## Abstract

Enterohemorrhagic *Escherichia coli* O157:H7 is an enteric pathogen responsible for bloody diarrhea, hemolytic uremic syndrome, and in severe cases even death. The study of O157:H7 is difficult due to the high specificity of the bacteria for the human intestine, along with our lack of sufficiently complex human cell culture models. The recent development of human intestinal enteroids derived from intestinal crypt multipotent stem cells has allowed us to construct 2-dimensional differentiated epithelial monolayers grown in transwells that mimic the human intestine. Unlike previous studies, saline was added to the apical surface, while maintaining culture media in the basolateral well. The monolayers continued to grow and differentiate with apical saline. Apical infection with O157:H7 or commensal *E. coli* resulted in robust bacterial growth from 10^5^ to over 10^8^ over 24 hours. Despite this robust bacterial growth, commensal *E. coli* neither adhered to nor damaged the epithelial barrier over 30 hours. However, O157:H7 was almost fully adhered (>90%) by 18 hours with epithelial damage observed by 30 hours. O157:H7 contains the locus of enterocyte effacement (LEE) pathogenicity island responsible for attachment and damage to the intestinal epithelium. Previous studies report the ability of nutrients such as biotin, D-serine, and L-fucose to downregulate LEE gene expression. O157:H7 treated with biotin or L-fucose, but not D-serine displayed both decreased attachment and reduced epithelial damage over 36 hours. These data illustrate enteroid monolayers can serve as a suitable model for the study of O157:H7 pathogenesis, and identification of potential therapeutics.

**Importance:** O157:H7 is difficult to study due to its high specificity for the human intestine and the lack of sufficiently complex human cell culture models. The recent development of human intestinal enteroids derived from intestinal crypt multipotent stem cells has allowed us to construct 2-dimensional differentiated epithelial monolayers grown in transwells that mimic the human intestine. Our data illustrates enteroid monolayers can serve as a suitable model for the study of O157:H7 pathogenesis, and allow for identification of potential therapeutics.

## Introduction

Enterohemorrhagic *Escherichia coli* O157:H7 is a food-borne pathogen typically acquired through contaminated food and water. Following consumption, O157:H7 travels to the distal end of the ileum. The bacteria directly attach to the intestinal epithelium and causes microvillus effacement. Activation of the locus of enterocyte effacement (LEE) pathogenicity island results in expression of a type-3 secretion system (T3SS), and injection of virulence genes into the cytoplasm of the intestinal epithelium (1, 2). One injected protein is the translocated intimin receptor (Tir), which binds a transmembrane protein on O157:H7 known as intimin (3). The Tir-intimin axis facilitates tight attachment and injection of additional virulence factors by the O157:H7 T3SS. A key non-LEE characteristic of O157:H7 is phage encoded Shiga Toxin (Stx) (4). Stx is a protein toxin that deadenylates the 28S rRNA in the 60S ribosomal subunit, inhibiting protein synthesis and eventually inducing cellular death (5, 6). Robust Stx expression during severe infection can result in hemolytic uremic syndrome, acute kidney damage, and even death. Stx is encoded by late phage genes and is only expressed during the lytic phase. Stx expression is controlled by a regulatory network under the bacterial SOS response (7). The bacterial SOS response can be activated by numerous stress responses present within the intestine. The most noteworthy being antibiotic exposure (8). Due to the activation of phage production and Stx expression during antibiotic exposure, there is a lack of viable treatment options for infection. Currently, only palliative treatment being used resulting in an urgency to further understand O157:H7 infection and identify potential therapeutics.

The study of O157:H7 is complicated by the high species specificity for the human intestine. This specificity makes common animal models such as mice and other rodents infeasible to use for studying attachment. Many studies have been performed in human transformed intestinal cell lines, which are not capable of fully recapitulating the human intestine. They do not produce the differentiated cell types present in the intestinal epithelium, and often contain multiple mutations. Recent developments in stem cell technology have led to breakthroughs in using Human Intestinal Organoids (HIOs) and Human Intestinal Enteroids (HIEs) as novel models for the study of various gastrointestinal pathogens (9–12). HIOs are produced from pluripotent stem cells, and when differentiated produce both a differentiated intestinal epithelium, and the surrounding mesenchymal cells (13, 14). HIOs respond to O157:H7 and act as a suitable model for studying infection (15). However, micro-injecting bacteria into the lumen is difficult. HIOs do not give adequate information regarding bacterial attachment to the epithelium making them inadequate for some studies. HIEs are derived from multipotent stem cells within intestinal crypts and produce only the intestinal epithelium. HIE can be growth both as 3D spheroids but can also be plated on transwell membranes for HIE monolayers (HIEMs). Once plated, HIEMs differentiate with a columnar structure containing basolateral nuclei and an apical actin surface as expected in the intestine. Previous studies with HIEMs typically utilize differentiation media in the apical transwell (12, 16). Differentiation media creates a nutrient rich environment for robust bacterial growth and can alter bacterial gene expression not typically seen within the intestinal lumen. To overcome these limitations, in this study we grew HIEs in transwells, forming a monolayer. We have replaced the apical differentiation medium with saline, and infected them with commensal, probiotic, and pathogenic *E. coli*. Our data demonstrates the utility of this HIEM system for the study of *E. coli* infection.

## Materials and Methods

### Growth of HIEMs

HIEs were derived from the H1 Line (WA01) NIH Registration #0043 human embryonic stem cell line with a normal 46, XY karyotype as previously described (Pradhan et. al., 2020). For development of the epithelial monolayers, transwell membranes (cat. 7430; Corning) were used, and cells were plated according to previously defined methods (Zuo et. al. 2018). Epithelial stem cells were grown in IntestiCult™ Organoid Growth Medium (Human) (cat. 06010; Stemcell) with 10 μM Y-27632 (cat. 72302: Stemcell) in both apical and basolateral chambers overnight. On day 2 the apical and basolateral media were replaced with differentiation media, containing a 1:1 solution of Intesticult OGM Human Basal Media (cat. 100-0190; Stemcell), and Gut Media containing DMEM/F12 (cat. DF-042-B; Millipore), B27 insulin (cat. 17504044: Invitrogen), N2 supplement (cat. 17502048: Invitrogen), 2 mM L-glutamine (cat. SH3003401; Fisher), 15 mM HEPES (cat. 15630080; Invitrogen), 100 ng/mL epidermal growth factor (cat. 236-EG-200; R&D Systems), and 2 mM penicillin/streptomycin (Pen/Strep) (cat. 15140-122; Invitrogen). Monolayers were grown until confluency (approximately 8 days) with apical and basolateral media changes every 2 days. Once confluent the apical differentiation media was replaced with sterile grade saline (cat. Z1376, Intermountain Life Sciences) while maintaining basolateral differentiation media. The basolateral media continued to be changed every 2 days, with saline only being changed once a week for 14 days post saline addition. Experiments were performed on day 14 post saline replacement.

### Bacterial strains, Growth of *E. coli*

The strains used in our study have been previously characterized, with assemblies and sequencing data for strains ECOR13 and PT29 deposited under NCBI BioProject ID: PRJNA 359210. *E. coli* O157:H7 PT29S was used for all O157:H7 infections. PT29S is a spontaneous streptomycin mutant of PT29 previously isolated from a patient, PT29S only contains genes for the more potent Shiga toxin 2 (Stx2), while lacking the genes for Stx1. PT29S also contains virulence genes associated with the Locus of Enterocyte Effacement (LEE) for attachment and effacement of the intestinal epithelium. ECOR13 is a non-pathogenic *E. coli* isolated from a healthy individual in Sweden. ECOR13 was obtained from the Michigan State University STEC Center for ECOR collection.

### HIEM Infections

O157:H7 was grown in Luira broth (LB) (cat. L24040, RPI) with or without biotin (100 nM), D-serine (100 μM) and L-fucose (100 μM) overnight prior to infection. On day zero of infection, optical density at 600 nm (OD_600_) was recorded with a Spec20D Spectrometer, using the conversion factor of OD 1.0 is equivalent to 1×10^9^ CFU/mL. Dilutions were performed to allow for an initial infection of 1×10^5^ bacteria within 100μL of saline.

In most studies immediately prior to infection, the monolayer media was removed and replaced with antibiotic-free medium. Infections were performed with an initial dose of 1×10^5^ bacteria in the apical saline, unless otherwise noted. Biotin, D-serine, and L-fucose were added at concentrations of 100 nM, 100 μM, and 100 μM respectively to both apical saline and basolateral media at time of initial infection. Transepithelial electrical resistance (TEER) was recorded immediately following addition of bacteria using the ERS-1 volt-ohm meter (Millicell). TEER was recorded over the course of the experiment, and upon sample collection. All apical and basolateral media was collected and plated for quantification of CFUs. Samples were either collected for confocal microscopy or lysed for quantification of bacterial adherence.

#### Determination of unattached and basolateral bacteria

Apical saline and basolateral media were collected. Serial dilutions were performed on the apical saline and 10 μL was plated on LB with 1.5% agar. Basolateral media was plated undiluted, unless bacteria were present, then serial dilutions were performed. CFUs were then counted and quantified as CFUs per well for each transwell. For the basolateral media 10 μL was plated out of the total 500 μLs, giving the basolateral bacteria a limit of detection (LOD) of 50 CFU/mL.

#### Determination of attached bacteria

Following saline removal, monolayers were washed with PBS three times. Epithelial cells were then lysed with 0.1% SDS for 10-15 minutes at 37°C. Following lysis, the lysate was collected into PBS bringing the total volume to 1 mL, allowing for CFU per mL to equal CFU per well for our attached bacteria. Serial dilutions were performed, and then 10 μL per dilution was plated followed by CFU counting, and quantification of CFUs per mL.

### Immunofluorescent (IF) Staining

Monolayers collected for IF, and confocal microscopy were washed with PBS three times to remove any unattached bacteria. Cells were fixed with 4% paraformaldehyde for 15 minutes at room temperature. Cells were washed with PBS and placed in 30% sucrose and stored at 4°C until staining could be performed. For staining, the 30% sucrose was removed, and cells were washed with three times with PBS with 0.1% Tween-20 (PBST). Monolayers were permeabilized with 0.2% Triton X-100 in PBS for 10 minutes at room temperature, then washed twice with PBST. Followed by blocking for two hours at room temperature in 10% goat serum, 0.1% BSA, and 0.01% Triton X-100 in PBS. Cells were washed with PBST and placed in primary antibody (Rabbit Anti-*E. coli*, cat. 1001, ViroStat, Rabbit Anti-Intimin, cat. NR-12194, BEI Resources, Mouse Anti-*E. coli* Serotype O157:H7, cat. MA5-18196, Invitrogen, Rabbit Anti-Villin, cat. Ab130751, Abcam) diluted in 1:200 in blocking buffer, as described above, at 4°C overnight. Primary antibody was removed the following day, and cells washed with PBST three times. Secondary AlexaFluor antibodies (488 Goat Anti-Mouse, cat. A11001, Life Technologies, 488 Goat Anti-Rabbit, cat. A32731, Invitrogen, 660 Goat Anti-Mouse, cat. A21055, Invitrogen, 660 Goat Anti-Rabbit, cat. A21074, Invitrogen) were added 1:500 for two hours in the dark at room temperature. Monolayers were washed with PBST three times and stained with Texas Red Phalloidin (cat. T7471, Invitrogen) diluted 5:200 in PBS for 30 minutes at room temperature in the dark. The cells were washed with PBST, followed by DAPI (cat. D3571, Invitrogen) incubated at room temperature for 5 minutes. Cells were washed twice with dH_2_O. A scalpel was used to circumvent the membrane and remove it from the transwell insert. The membrane was placed on a microscope slide with the cells facing upwards away from the slide, and vectamount permanent mounting medium (cat. H-5000-60; Vector) was used to mount the coverslip. Mounting medium was allowed to dry overnight, and coverslips were circumvented with clear nail polish to prevent movement during imaging. Stained sections were imaged using a Zeiss LSM710 Live Duo Confocal Microscope (University of Cincinnati Live Microscopy Core).

### Quantification and Statistical Analysis

All statistical analysis was done using Graphpad Prism 5. Statistical tests used are indicated in figure legends.

## Results

### HIEMs Can Grow with Apical Saline

HIEMs were plated in transwells in differentiation medium. The apical medium was replaced with saline 7-8 days later, once the monolayer was fully confluent, and media could not freely transfer between the basolateral and apical media layers. The monolayers were cultured for 14 days after saline addition. The basolateral differentiation media was changed every 2-3 days. TEER was recorded at the time of every media change and compared with monolayers with apical media (Figure 1A). The TEER was similar in the medium and saline wells up until day 14, when the TEER for the medium continued to rise while the TEER for the saline plateaued. This is most likely due to the apical media facilitating multiple layers of cells due to the abundance of nutrients, while this phenomenon was avoided with apical saline replacement. The monolayers were characterized by confocal microscopy. Fully confluent monolayers with a single layer of cells had formed, containing cellular junctions following 14 days of saline addition (Figure 1B).

**Figure 1.**
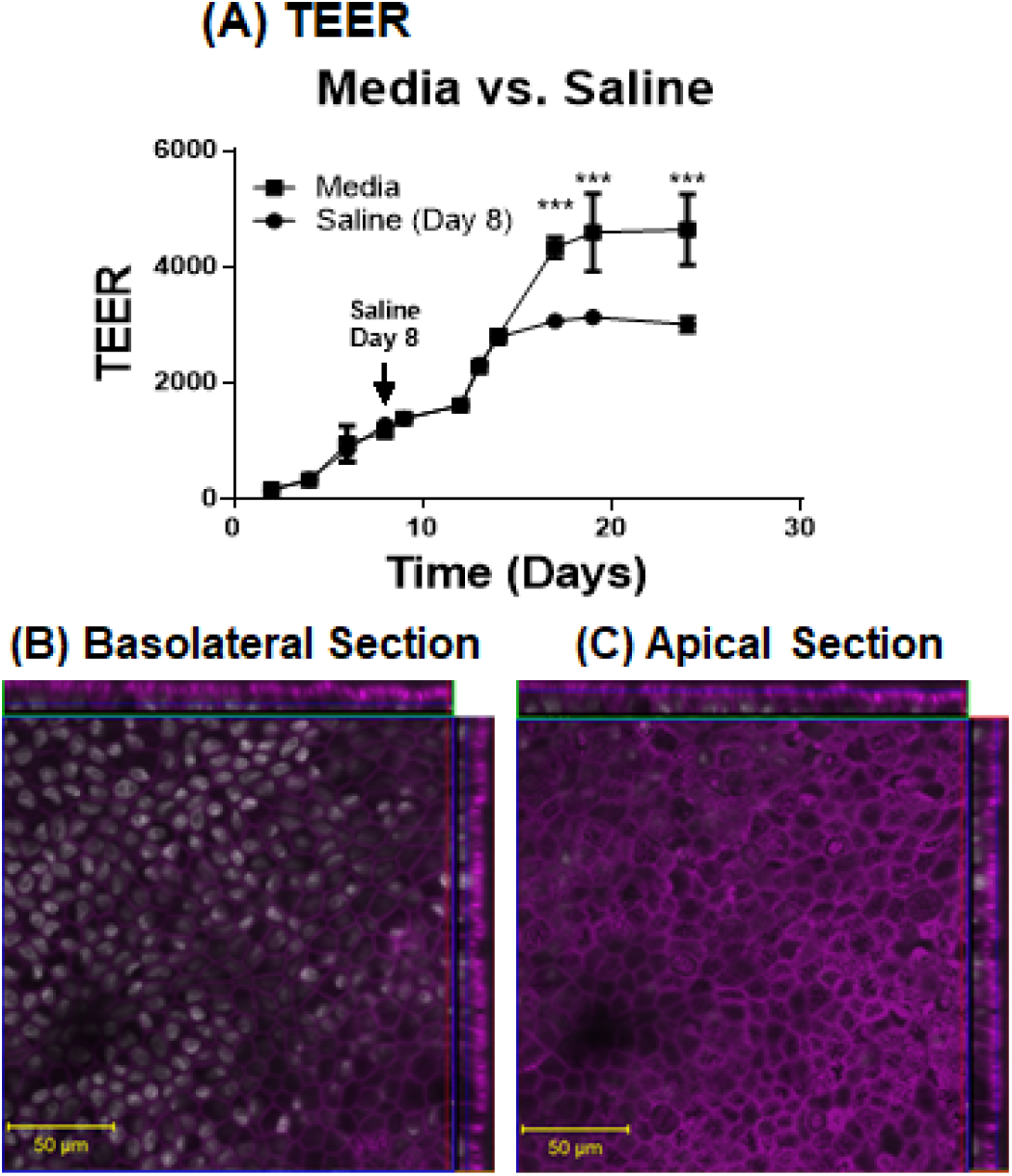
HIEMs grow and differentiate following apical saline replacement. Apical differentiation media was replaced with saline on day 8 and grown for an additional 14 days with only basolateral media changes. A) TEER continued to increase to a maximum of 2500. B) Confocal microscopy stained with DAPI (white), and actin (magenta). All data was collected in triplicate, and statistical analysis was done using a two-way ANOVA with Sidak’s multiple comparisons test.

These data indicate our monolayers can grow, properly polarize, and maintain a confluent 2D monolayer within the transwells.

### HIEMs Response to Infection with Commensal & Probiotic E. coli

Initial infection of 10^4^ with probiotic *E. coli* Nissle 1917 was performed, with and without basolateral Pen/Strep **(Figure 2A)**. TEER was recorded immediately post infection, and 24 hours post infection. TEER was maintained over the course of infection with or without Nissle **(Figure 2B)**. Monolayers were harvested for both CFUs and IF at 24 hours post infection. CFUs indicated robust bacterial proliferation increasing to greater than 10^6^ **(Figure 2C)**. There was no basolateral growth over 24 hours, regardless of pen/strep addition (data not shown). Confocal microscopy was performed. Z-Stack images were collected from Nissle infected monolayers at 24 hours post infection and stained for nuclei (white), actin (magenta), and Nissle (green). A fully confluent monolayer with basolateral nuclei was observed **(Figure 2D)**. A confluent actin surface with small numbers of bacteria **(Figure 2E, green)** associated with the monolayer was seen in the apical view. Epithelial polarization is confirmed additionally with the Z view on the side of each image illustrating the localization of the nuclei, and actin barrier.

**Figure 2.**
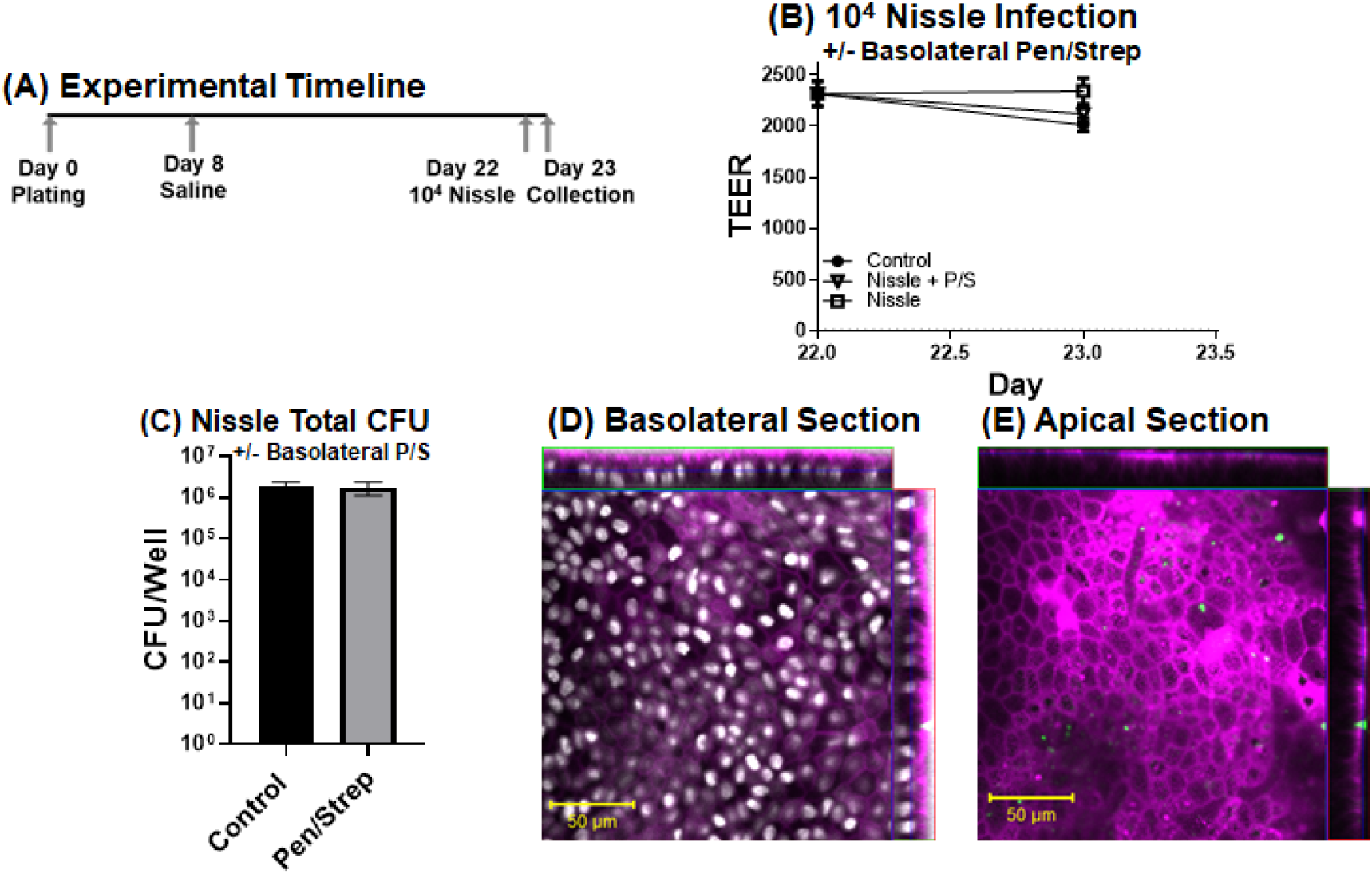
Infection with Nissle 1917 shows no loss in epithelial barrier integrity over 24 hours. **A)** Experimental outline. HIEs were plated, apical media replaced with saline on day 8, and infection with 10^4^ Nissle was preformed 14 days post saline replacement. Samples were collected 24 hours post infection. **B**) HIEMs were infected with Nissle in the presence or absence of basolateral antibiotics. TEER was not changed over 24 hours post infection. **C**) CFUs were quantified for total bacteria growth with and without the presence of basolateral antibiotics. **D-E**) Confocal microscopy of Nissle infected HIEMs. **D**) basolateral and **E**) apical section stained for DAPI (white), actin (magenta), and *E. coli* (green).

The infection was increased to 10^5^ using both commensal *E. coli* ECOR13, and probiotic Nissle 1917. Similar results were seen at 24 hours; there was no loss of epithelial integrity shown by TEER **(Figure 3A)**. Assessment of bacterial growth showed robust bacterial proliferation to 10^8^, with minor bacterial attachment (<5%). No growth of basolateral bacteria was seen for ECOR13 **(Figure 3B)**, while one of the three cultures of Nissle was positive **(Figure 3C)**. The rapid replication of the bacteria is not fully understood, but it is likely the bacteria are using intestinal mucus as the primary carbon source for replication.

**Figure 3.**
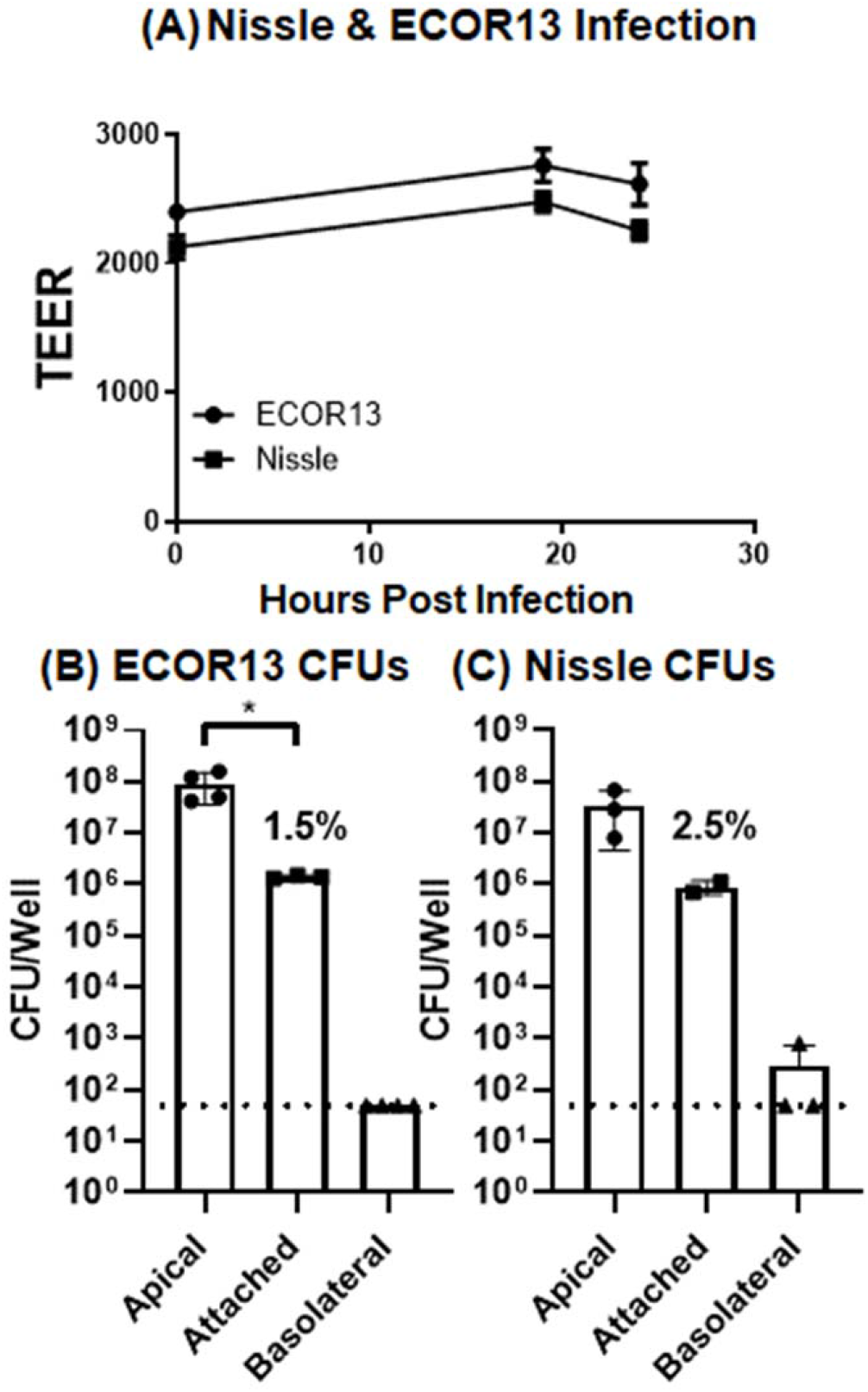
HIEM infections with 10^5^ of probiotic Nissle or commensal *E. coli*. HIEMs were infected with 10^5^ Nissle or ECOR13 (commensal *E. coli*). **A)** TEER values were similar over 24 hours of infection. **B–C)** CFU showed apical growth from 10^5^ to >10^7^, low epithelial attachment, and low basolateral bacteria. **B)** ECOR13 and **C)** Nissle. The dotted line represents the limit of detection (LOD) of 50 CFUs per well. Statistical analysis was done using an unpaired t-test between apical and attached bacteria with * p<0.05.

These data indicate the ability of monolayers to retain their barrier integrity following exposure to commensal and probiotic bacteria. The non-pathogenic bacteria gain nutrients from the monolayers for rapid replication, without damaging the epithelium in the process, and without attaching in high quantities.

### HIEMs Response to Infection with O157:H7

HIEMs were infected with O157:H7 at a dose of 10^5^ bacteria, and TEER was quantified over 36 hours of infection **(Figure 4A)**. TEER remained constant for 12 hours, then dropped, a trend that continued until 36 hours post infection where the TEER was decreased to 1000 Ohms. Quantification of CFUs was done at multiple time points over the course of infection **(Figure 4B)**. Bacteria replicated from 10^5^ to 10^8^ over 36 hours of infection and reached a peak at approximately 24 hours. *E. coli* O157:H7 was almost fully adhered (>90%) by 24 hours post infection. This is also when TEER began to drop, indicative of epithelial damage. The presence of basolateral bacteria was quantified. Basolateral bacteria growth was detected by 30 hours post infection and continued to increase through 36 hours post infection, illustrating epithelial damage and loss of barrier integrity allowing for bacterial translocation between apical and basolateral chambers.

**Figure 4.**
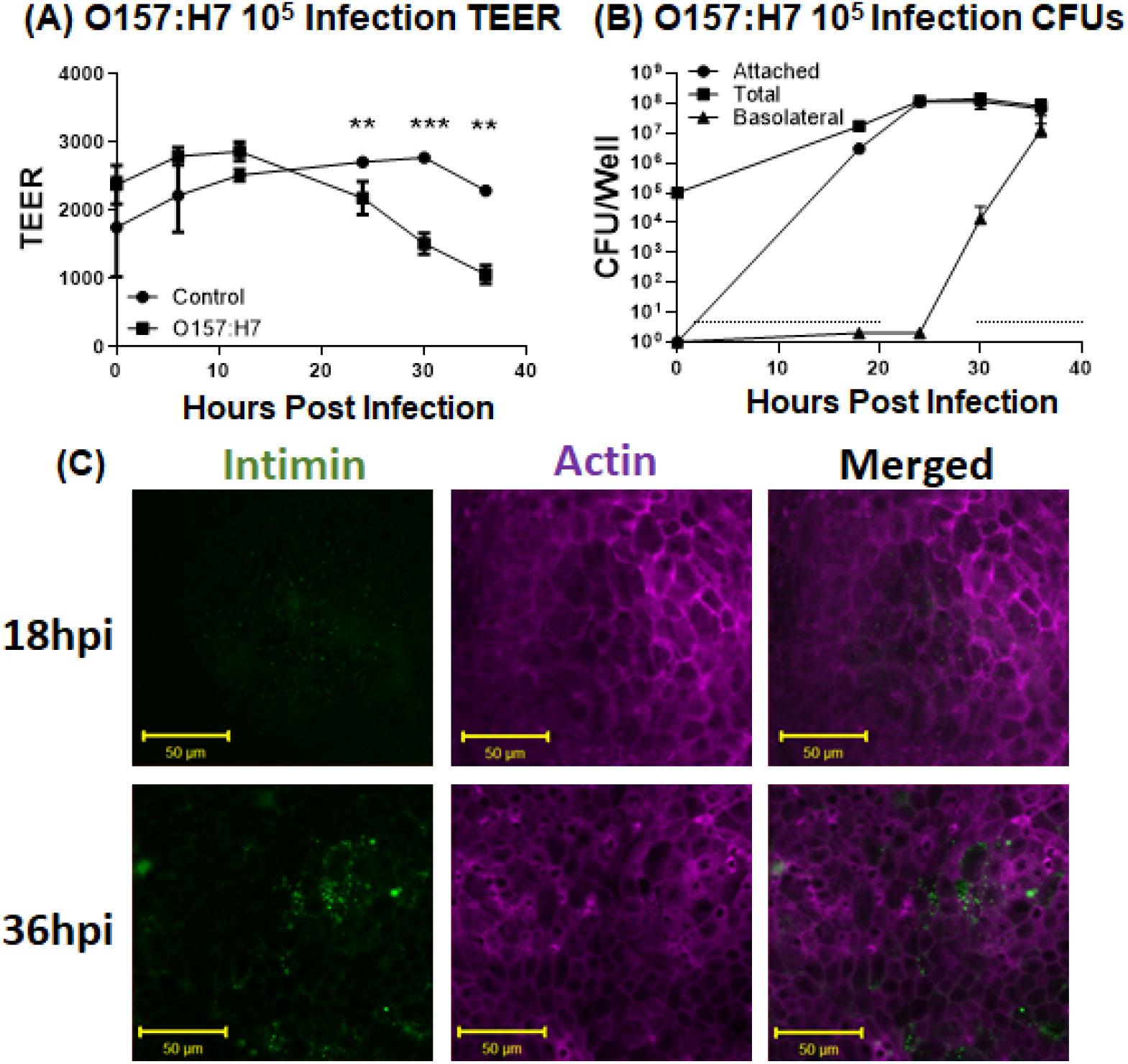
O157:H7 damaged the epithelial monolayers. HIEMs were infected with 10^5^ O157:H7. **A)** TEER was recorded for 36 hours post infection. TEER was reduced by 24 hours post infection, and beyond using a mixed-effect analysis, p<0.05 *, P<0.005 **, P<0.001 ***. **B)** Total bacteria increased from 10^5^ to 10^8^ over 24 hours of infection. Basolateral bacteria were detected at 30 hours post infection and increased through 36 hours, indicative of epithelial damage. Attached bacteria reached levels of >95% attached by 24 hours post infection and remained constant to 36 hours post infection. **C)** HIEMs were stained at 18- and 36-hours post infection for actin (magenta) and intimin (green).

To assess the expression of intimin by O157:H7, confocal microscopy was performed on epithelial cells at 18 and 36 hours post infection **(Figure 4C)**. At 18 hours post infection intimin can be seen in small sections of the epithelial surface **(Figure 4C, top)**. By 36 hours post infection a sharp increase in the amount of intimin can be seen as compared to 18 hours post infection **(Figure 4C, bottom)**. Assessment of monolayers 24 hours post infection illustrates epithelial damage **(Figure 5)**, as indicated by extrusion of epithelial nuclei **(Figure 5B, arrow)**.

**Figure 5.**
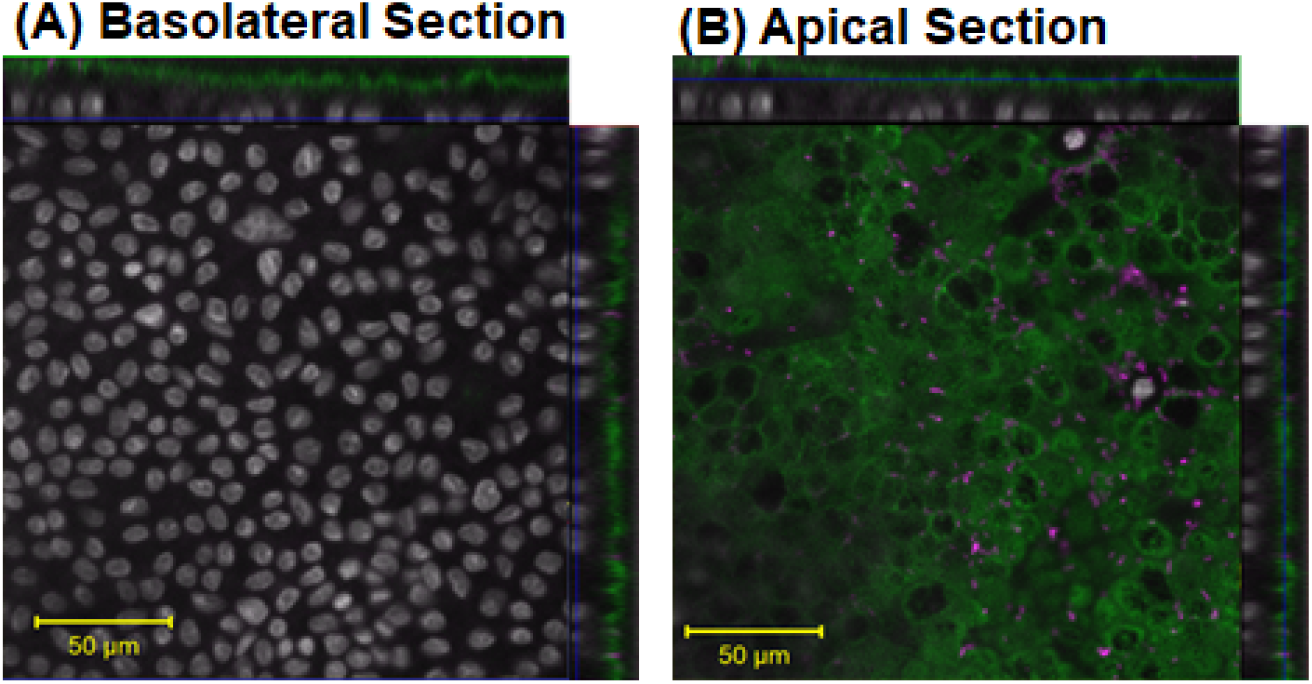
O157:H7 induces epithelial damage following attachment. HIEMs infected with 10^5^ O157:H7 were collected 24 hours post infection and stained for villin (green), O157:H7 (magenta), and DAPI (gray). Z-stacks were taken, and A) basolateral, and B) apical section are shown. Extruding nuclei were seen (arrow).

These data demonstrate that O157:H7 directly attaches to the epithelium and induces epithelial damage to the HIEM. Intimin staining confirmed O157:H7 utilizes the Tir-intimin mechanism of attachment facilitated through the T3SS. Following epithelial damage HIEMs exhibit the ability to extrude damaged epithelial cells from the lumen.

### Rescue with biotin, L-fucose

Expression of the O157:H7 T3SS, intimin, and downstream virulence genes are all expressed in the Locus of Enterocyte Effacement (LEE) pathogenicity island. LEE genes are regulated by multiple signals present within the intestine that vary through the length of the gastrointestinal tract. This regulatory network allows the bacteria to express the LEE genes at the distal end of the ileum, the preferred attachment location. Three factors, biotin (17), fucose (18), and D-serine (19) **(Figure 6A)** have been reported to decrease LEE gene expression and reduce epithelial attachment. L-fucose acts via FusKR, biotin acts via Fur, and D-serine acts via GadE transcription factors to prevent LEE gene expression.

**Figure 6.**
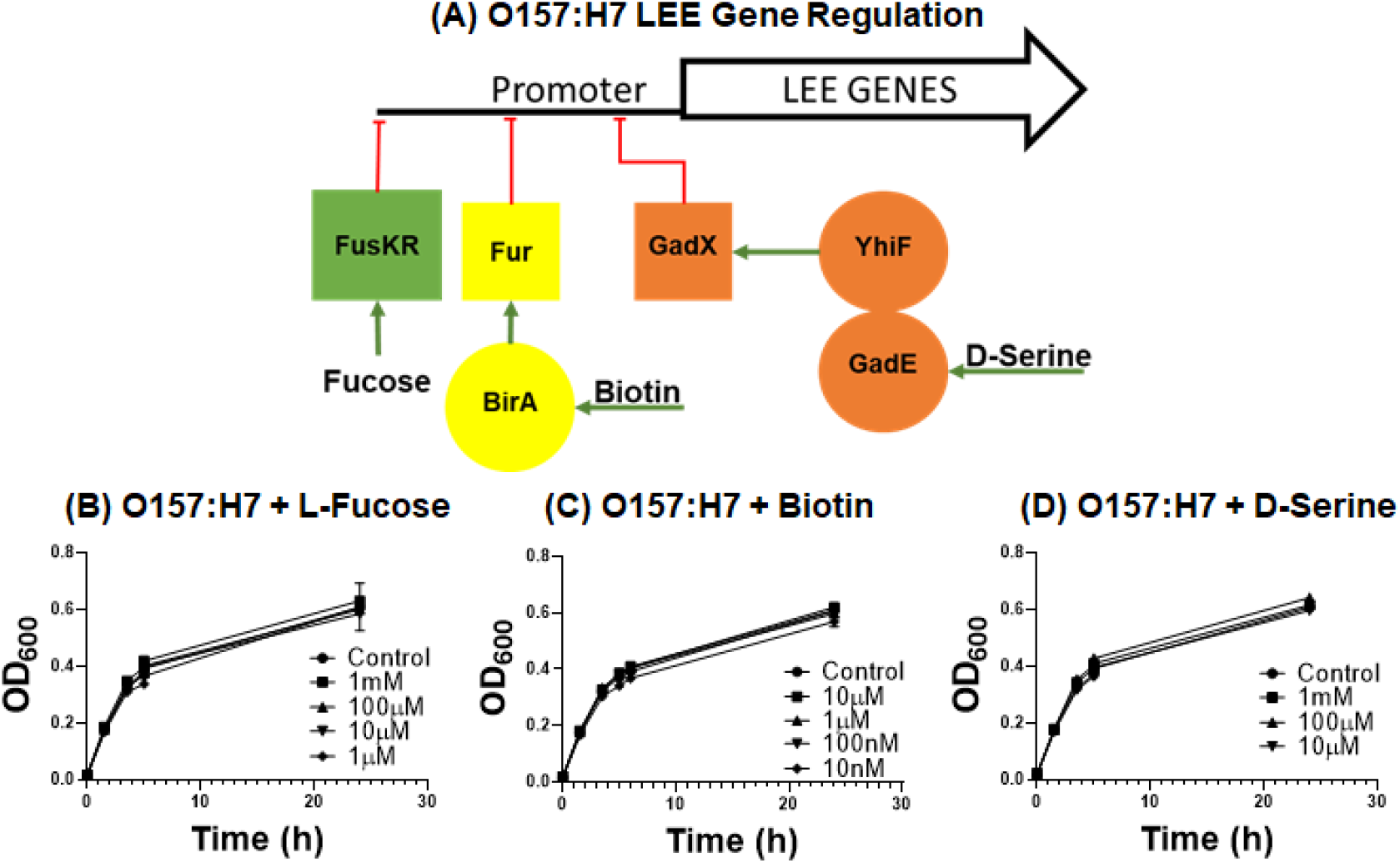
Schematic of O157:H7 LEE gene expression. A) L-fucose acts via FusKR, biotin acts via BirA, and D-Serine acts via GadE, preventing LEE gene expression. (B) fucose, (C), biotin, (D) nor D-serine, do not inhibit O157:H7 growth in LB over 24 hours.

Initial experiments were performed by growing O157:H7 in LB containing increasing concentrations of L-fucose **(Figure 6B)**, biotin **(Figure 6C)**, and D-serine **(Figure 6D)**. No impact on bacterial growth was seen over 24 hours. We then use these cultures to study O157:H7 infection. Compared to untreated infections, O157:H7 treated with 100 μM biotin prior to and during monolayer infection maintained TEER levels **(Figure 7A, top)**. Similar results were seen with L-fucose **(Figure 7B, top)**. Reduced epithelial attachment was also observed in biotin **(Figure 7A, bottom)**, and L-fucose **(Figure 7B, bottom)** treated O157:H7 infections. The same experiment was performed with D-serine treated O157:H7, and no significant difference was seen in TEER or attachment (data not shown). These data correlate with previous studies showing a decreased epithelial attachment following O157:H7 treatment with biotin and L-fucose (17, 18).

**Figure 7.**
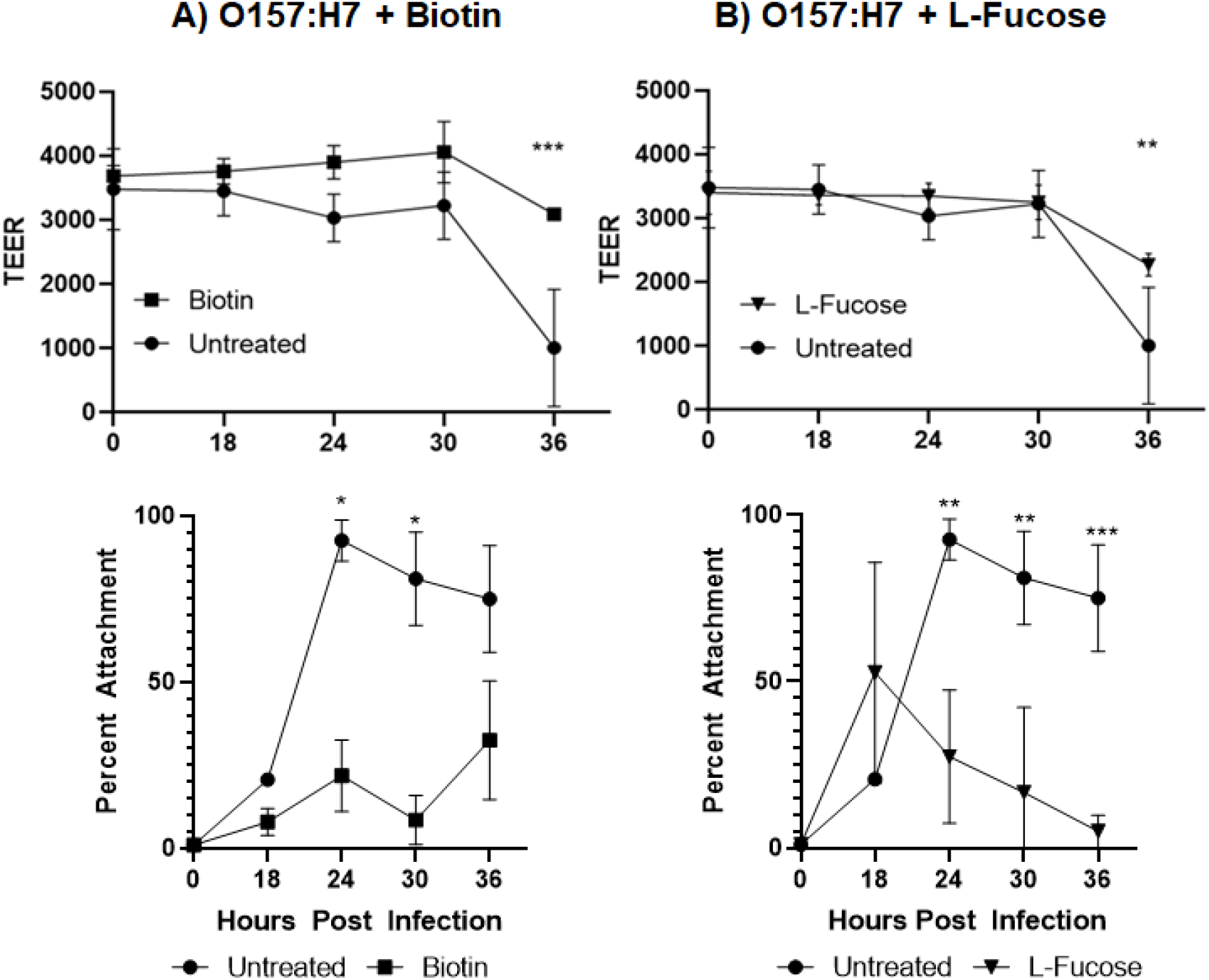
O157:H7 infection is limited by biotin and L-fucose over 36 hours of infection. HIEMs were infected with 10^5^ O157:H7 grown overnight in LB containing 100 nM biotin, 100 μM L-fucose, or control cells. **A)** Treatment of O157:H7 with biotin prevented loss of TEER (**A, top**) and reduced epithelial attachment (**A, bottom**). **B)** Similar results were seen with L-fucose for both TEER (**B, top**) and attachment (**B, bottom**). Statistical analysis was done using mixed-effect analysis with p<0.05 *, P<0.005 **, P<0.001 ***.

## Discussion

This study focuses on utilizing recent developments in stem cell technology to create intestinal epithelial monolayers that properly mimic intestinal physiology, and host-pathogen interactions. The presence of apical medium is not consistent with the luminal environment. The lumen is not buffered and presence of glucose in the medium allows for bacterial growth and generation of acid. To create an environment more consistent with the lumen, we replaced the medium with saline. We demonstrate that apical medium is not needed for differentiation of the epithelial monolayers or to maintain proper polarization. Furthermore, commensal, probiotic and pathogenic *E. coli* rapidly grew in this environment. Despite being suspended in saline, the bacteria were able to grow over the course of infection, increasing by over 3 logs. It is not entirely clear what energy source is utilized by the bacteria for robust growth, but it is hypothesized mucus secreted by the epithelium is degraded by *E. coli*, and acts as the primary carbon source in the human intestine (20, 21)

The goal of our study was to focus on the interaction between O157:H7 and the intestinal epithelium. Infection with both commensal and probiotic *E. coli* strains did not induce epithelial damage and the bacteria did not attach to the apical surface. O157:H7 also underwent robust bacterial growth; however, greater than 90% of the bacteria were directly attached to the intestinal epithelium. Following attachment, the epithelium was damaged. Loss of barrier integrity started at 24 hours post infection. Based on immunofluorescence and microscopy for intimin this process is facilitated through the expression of LEE genes, and utilization of the Tir-intimin attachment system.

Both biotin and L-fucose reduced attachment of O157:H7 to the epithelium, resulting in less epithelial damage later in infection, as evidenced by maintenance of TEER. There are currently no treatments for O157:H7 infection, and it is tempting to suggest that these simple nutrients can be used therapeutically.

Our data illustrates the utility of human intestinal enteroid monolayers with apical saline replacement as a model to study O157:H7, and potentially other intestinal pathogens. This work can begin to fill a gap for a physiologically relevant model for the study of O157:H7, and the identification of potential therapeutics.

## Current Funding

U19 – AI116491, R01 – AI139027

NIDDK P30 DK078392 – Pluripotent Stem Cell and Organoid Core and Live Microscopy Core of the DHC.

